# The stress response factor SigH mediates intrinsic resistance to multiple antibiotics in *Mycobacterium abscessus*

**DOI:** 10.1101/2025.03.04.641476

**Authors:** Md Shah Alam, Mst Sumaia Khatun, Buhari Yusuf, Lijie Li, Aweke Mulu Belachew, Haftay Abraha Tadesse, Jingran Zhang, Xirong Tian, Cuiting Fang, Yamin Gao, Zhiyong Liu, H.M. Adnan Hameed, Jinxing Hu, Xinwen Chen, Nanshan Zhong, Shuai Wang, Tianyu Zhang

## Abstract

*Mycobacterium abscessus* (Mab) causes pulmonary diseases with limited treatment options due to its high level of intrinsic resistance to available drugs. Mab possesses complex and poorly understood drug resistance mechanisms. Identifying new drug targets and gaining a deeper understanding of drug resistance mechanisms are essential for discovering novel therapeutic alternatives. Here, we investigated the role of a putative sigma factor SigH in intrinsic multi-drug resistance in Mab. Mab SigH shares an 84% peptide sequence identity with *Mycobacterium tuberculosis* (Mtb) SigH, a well-known stress response protein and global transcriptional regulator. We constructed a *sigH* gene deletion strain of Mab (Δ*sigH*) and complemented strains by expressing either Mab *sigH* (CPMab*sigH*) or Mtb *sigH* (CPMtb*sigH*) in Δ*sigH.* The Δ*sigH* strain exhibited hypersensitivity to a broad range of antibiotics, including levofloxacin, moxifloxacin, tigecycline, tetracycline, amikacin, vancomycin, and rifabutin and all complemented strains restored the drug resistance phenotype. Additionally, Δ*sigH* showed increased sensitivity to oxidative and heat stress compared to the wild-type Mab and complemented strains. Transcriptomic analysis revealed that deletion of *sigH* disrupted the balance of gene expression, primarily elevating the expression of genes encoding YrbE and MCE family proteins and downregulating genes expressing ABC-type transporters, sigma and anti-sigma factors and other genes associated with antimicrobial resistance. Collectively, our findings indicate that SigH is a key regulator of global gene expression in response to environmental stresses, including antimicrobial treatment, and is crucial for the intrinsic drug resistance of Mab. SigH represents a promising target for the development of novel therapeutic strategies against Mab infections.

## Introduction

*Mycobacterium abscessus* complex (MABC) includes well-known rapidly growing non-tuberculosis mycobacteria (NTM) capable of causing severe acute and chronic lung infections, particularly in patients with underlying lung diseases such as cystic fibrosis and obstructive pulmonary diseases, as well as skin and soft tissue infections (1, 2). Recently, MABC has been associated with a wide range of clinical manifestations due to increasing global morbidity and mortality rates (3). The intrinsic drug resistance mechanisms of MABC remain poorly understood, and the highly resistant phenotype of *Mycobacterium abscessus* (Mab) presents a significant challenge in its infection therapy. To improve treatment options and combat prevalent Mab infections, novel drugs with new mechanisms of action are urgently needed.

MABC accounts for 2.6-13.0% of all NTM-related pulmonary infections (4). Mab demonstrates intrinsic resistance to most therapeutic agents, and cure rates for Mab lung infections are very low (approximately 25-58%), earning it the moniker “antibiotic nightmare” (5). The primary mechanisms of intrinsic drug resistance include a waxy, impermeable cell wall, drug efflux pump systems, and drug inactivation by hydrolases or modifying enzymes (6, 7, 8). For instance, aminoglycoside phosphotransferases and 2′-N-acetyltransferases transfer acetyl or phosphate residues to specific positions within aminoglycoside drugs, rendering them inactive (9). Erythromycin resistance methylase is responsible for macrolide resistance, while MmpL5/MmpS5 confers resistance to bedaquiline and clofazimine (10). The MabTetX, a WhiB7-independent tetracycline-inactivating monooxygenase, increases Mab’s resistance to the tetracycline family (11). Additionally, spontaneous mutations in particular genes in response to drugs cause acquired resistance. For example, genetic polymorphism results in resistance to antibiotics like fluoroquinolones (FQs), which are broad-spectrum secondary therapeutic agents for multi-drug-resistant tuberculosis and act by inhibiting DNA gyrase supercoiling activity (12, 13). Mab resistance to FQs is due to genetic alterations, especially in quinolone resistance-determining regions within DNA gyrase subunits GyrA and GyrB, the primary targets of FQs (14). Tigecycline (TIG) resistance in Mab has been linked to both SigH and RshA (15). Dysregulated environmental stress response sigma factor (SigH) is associated with tigecycline resistance in both *Mycobacterium tuberculosis* (Mtb) and Mab (16, 17). The limited success in anti-Mab drug discovery primarily stems from high intrinsic resistance and rapidly acquired resistance to currently available active drugs. However, the mechanisms underlying intrinsic drug resistance in Mab remain not fully elucidated (18).

WhiB7 is a redox-sensitive master transcriptional regulator crucial for activating intrinsic drug resistance systems in mycobacteria and other bacteria, such as *Streptomyces lividans* and *Rhodococcus jostii* (19, 20). The multi-drug resistance WhiB7 transcriptional regulator induces SigH transcription. SigH is an alternative sigma factor involved in the transcriptional regulation of genes responsible for mycobacterial stress responses, including oxidative, heat, and nitrosative stresses (21, 22, 23), and is negatively regulated by RshA (15, 24). The significance of stress responses in etiology and immunity has been extensively studied in Mtb and *Mycobacterium smegmatis* (Msm) (22, 25).

In this study, we investigated the role of SigH (MAB_3543c) in antimicrobial resistance, stress response, and differential gene expression in Mab. Our findings lay the groundwork for further research that could lead to the development of new therapeutics or treatment regimens for Mab infections.

## Material and Methods

### Bacterial strains and growth conditions

The Mab GZ002 (accession number CP034181), exhibiting a smooth phenotype (26), was cultivated at 37 °C with shaking at 220 rpm. Growth media included Middlebrook 7H9 (Difco), supplemented with 0.2% glycerol, 0.05% Tween 80, and 10% OADC, or solid 7H10/7H11 agar supplemented with 0.5% glycerol and 10% OADC. *Escherichia coli* strain DH5α was propagated on solid or in liquid Luria-Bertani medium under identical conditions of 37 °C and 220 rpm shaking, with an incubation period of 10-12 hours, while Mab GZ002 was grown on agar medium for 3-5 days. Antibiotic usage and their respective concentrations were tailored to the specific experimental demands.

### CRISPR-Cpf1-assisted knockout of *sigH* (*MAB_3543c*)

CRISPR-Cpf1-assisted recombineering was employed to knockout the Mab *sigH* gene via homology-directed repair (27, 28, 29). Specifically, the CRISPR RNA (*crsigH*) was designed as two complementary oligonucleotides, each 24 base pairs in length (*crsigH*-F/R), targeting a region adjacent to a protospacer adjacent motif (PAM) characterized by the 5’-YTN-3’ trinucleotide sequence. One oligonucleotide was designed with a *Hin*dIII overhang, while the other had a *Bpm*I overhang. These oligonucleotides were annealed and cloned into *Hin*dIII/*Bpm*I- digested pCRZEO to generate pCRZEO-*crsigH*.

The homology-directed repair template (*sigH*UD) was constructed by amplifying 640-bp upstream (U) and 609-bp downstream (D) regions of the target gene, including flanking sequences from adjacent genes (Figure S2). Both fragments were cloned into pBlueSK to generate pBlueSK-*sigH*UD. *sigH*UD was amplified from pBlueSK-*sigH*UD using the primer pair *sigH*UD-F/R (Table S3). pCRZEO-cr*sigH* and *sigH*UD were transformed by electroporation into electrocompetent Mab cells harboring pJV53-Cpf1. The transformants were then plated on a 7H11 plate containing zeocin (ZEO, 30 µg/mL), kanamycin (KAN, 100 µg/mL), and anhydrotetracycline (ATc, 100 ng/mL), and incubated at 30 ^◦^C for 5 days. *sigH* knockout was verified by PCR and sequencing using the primer pair Id3543c-F/R (Figure S2). To construct the selectable marker-free knockout strain, the knockout strain was grown in 7H9 medium and plated on a drug-free 7H10 plate to obtain single colonies. A colony that grew on drug-free plates but not on KAN- and ZEO-containing plates was selected as the unmarked strain (Δ*sigH*) to be used in downstream experiments (Figure S3).

### Complementation and overexpression of *sigH*

Complemented (CP) strains were constructed by integrating *sigH* into the genome of Δ*sigH* to express the gene under its native promoter (*Np*) or *hsp60* promoter, or through ectopic expression – i.e. cloning *sigH* or its Mtb homolog into pMV261 and transforming them into Δ*sigH*. For overexpression, pMV261-*sigH* was transformed into wild-type Mab (WT). The transformants were grown on Middlebrook 7H10 agar plates supplemented with KAN at 100 µg/mL for 3 to 5 days at 37 °C and verified by PCR and sequencing. Thus, the following strains were used in this study: OEMab*sigH* (WT overexpressing Mab *sigH*), Δ*sigH* (selectable marker-free knockout strain for *sigH*), CPMab*sigH* (the CP strain ectopically expressing Mab *sigH*), CPMtb*sigH* (the CP strain ectopically expressing Mtb *sigH*), CP*Np*Mab*sigH* (the CP strain in which *Np*-Mab *sigH* is integrated into Δ*sigH* genome), and CP*hsp60*Mab*sigH* (the CP strain in which *hsp60*-Mab *sigH* is integrated into Δ*sigH* genome). Results of Mab strain verification and all primer sequences are provided in the supplementary material (Figures S4, S5; Table S3).

### Drug susceptibility testing

Mab strains were routinely cultivated on Middlebrook 7H10 agar or in Middlebrook 7H9 broth. Antimicrobial agents including TIG, tetracycline (TET), clarithromycin (CLA), clofazimine (CLF), vancomycin (VAN), amikacin (AMK), levofloxacin (LFX), moxifloxacin (MFX), imipenem (IMP), cefoxitin (CFX), rifabutin (RIB), and linezolid (LZD) were prepared as stock solutions and stored at −20 °C. Broth microdilution assay, adhering to Clinical Laboratory Standards Institute (CLSI) guidelines, was employed to determine drug susceptibility (30, 31). Mycobacterial cultures were standardized to approximately 1 × 10^7^ colony-forming units per milliliter (CFU/mL) in 7H9 medium without Tween 80. Bacterial suspensions underwent two-fold serial dilutions in the presence of individual drugs within 96-well plates. These plates were incubated at 37 °C for 3 days, with an extended incubation period of 14 days specifically for CLA, before assessing the endpoint. Minimum inhibitory concentrations (MICs) were established according to CLSI criteria, defining them as the minimal drug concentrations capable of visually inhibiting mycobacterial growth. Additionally, spot culture agar methodology was also utilized for drug susceptibility assessments (32). Herein, WT, Δ*sigH*, and CPMab*sigH* strains were propagated in 7H9 medium at 37°C until reaching an optical density (OD_600_ nm) of 0.6. Subsequently, ten-fold serial dilutions were applied and aliquoted onto plain Middlebrook 7H10 agar (Drug-free serving as a control) alongside plates supplemented with varying drug concentrations. Following a 3-day incubation at 37°C, the plates were examined.

### Thiol-specific oxidative and heat stress assay

WT, Δ*sigH*, and CPMab*sigH* were grown to the exponential phase and equilibrated at an OD_600_ nm of 0.6 for oxidative and heat stress assays. For oxidative stress assay, a 100 µL bacterial inoculum at OD_600_ nm 0.6 was spread on agar plates. Whatman paper disks (4.5 mm) were loaded with 10 µL oxidizing agent diamide at concentrations 4 M, 2 M, 1 M, and 0.5 M. These disks were then placed on agar plates and incubated for 3 days at 37 °C. The susceptibility of bacteria to diamide was assessed by measuring the zone of inhibition. Different survival curves were observed under heat and oxidative stress conditions. Survival under heat stress was determined based on CFU counts. Bacterial cultures were incubated in a water bath at 45 °C, and at 1-hour intervals, 100 µL of the bacterial culture was diluted in PBS and plated on 7H10 agar plates to determine the viable cell number (33). Additionally, bacterial survival was assessed in the presence of H_2_O_2_ and diamide. The bacterial OD_600_ nm was adjusted to 0.5-0.6 and cultured with 50 mM diamide and 50 mM H_2_O_2_ individually, followed by incubation at 37 °C. At one-hour intervals, bacteria were 10-fold subjected to serial dilutions and plated to count CFU for determining the survival rates under stress conditions (22, 34).

### RNA preparation, sequencing, and transcriptomic analysis

RNA isolation and library preparation for RNA sequencing were performed on the WT, Δ*sigH*, and CPMab*sigH* strains. These bacterial strains were cultured in Middlebrook 7H9 medium supplemented with Tween 80 and incubated at 37°C until reaching the exponential growth phase, characterized by an OD_600_ nm of approximately 0.6–0.8. RNA extraction was carried out using the TRIzol method, and sample quality was inspected using a Thermo NanoDrop One and Agilent 4200 Tape Station (35). Approximately, RNA samples were treated with the Epicentre Ribo-Zero rRNA Removal Kit (Illumina) to enrich for mRNA. Sequencing libraries were constructed using the NEBNext Ultra II Directional RNA Library Prep Kit. The constructed libraries underwent a quality inspection before being sequenced on Illumina’s high-throughput sequencing platform with PE150 configuration. The resulting reads were trimmed using fastp v0.23.2 (36) and mapped to the Mab reference genome (NCBI accession number CP034181). Quantitative analysis of gene expression levels was conducted to analyze the differentially expressed genes (DEGs) between different samples and to reveal the regulatory mechanisms of these genes by combining sequence function information. Gene expression was quantified using RNA sequencing by expectation maximization (RSEM), and fragments per kilobase of transcript per million mapped reads (FPKM) were calculated using Htseq-count (v0.11.2) (37, 38). Differential gene expression analysis was performed using DESeq2 and edgeR (39, 40), with the default screening conditions set to FDR ≤ 0.05 and |log_2_FC (FoldChange)| ≥ 1, applying Benjamini/Hochberg correction for multiple testing.

Functional annotation of the differential gene set was conducted using Gene Ontology (GO) to understand the roles of these genes, metabolic pathways, and other biological processes (41). Additionally, the Kyoto Encyclopedia of Genes and Genomes (KEGG) was used to determine the molecular functional pathways associated with DEGs (42). Functional enrichment analyses, such as GO enrichment and KEGG enrichment, were performed using cluster Profiler (43). Furthermore, Rockhopper software was employed for small RNA and transcript structure analysis (44).

### Ethidium bromide accumulation assay

The Ethidium bromide (EtBr) accumulation assay was conducted as previously described (45) to evaluate the cell envelope permeability of mycobacterial strains. Mycobacterial cultures were grown in Middlebrook 7H9 medium at 37 °C until reaching the mid-log phase. Bacterial suspensions were then normalized to an OD_600_ nm of 0.8 in phosphate-buffered saline (PBS) supplemented with 0.8% glucose. EtBr was added to the wells at a final concentration of 2 μ g/mL along with 0.4% glucose. Fluorescence measurements were taken using a Flex Station 3 Multi-Mode Microplate Reader (Molecular Devices, CA, USA), with excitation and emission wavelengths set at 530 nm and 590 nm, respectively. The fluorescence data from EtBr accumulation were recorded at 60-second intervals over a period of 60 minutes at 37°C. Data analysis and plotting were performed using GraphPad Prism version 10.3.1 (GraphPad, San Diego, USA).

## Results

### Identification of stresses response factor SigH in Mab

Mab is an opportunistic pathogen that causes morbidity in the presence of underlying conditions, including cystic fibrosis (CF). Condition-specific transcriptomics unveiled the molecular factors that drive the persistence and adaptation of Mab in its host. This was demonstrated through Mab exposure to synthetic CF sputum medium, which has been demonstrated to predominantly trigger strong up-regulation in the expression of sigma and anti-sigma factors (46). In addition, the role of these factors in stress response in bacteria has also been demonstrated (25). This therefore demonstrates their potential significance in Mab pathogenesis. These findings inspired us to investigate whether sigma and anti-sigma factors play a role in resistance to multiple antibiotics in Mab. Thus, we identified two adjacent sigma and anti-sigma factors, *sigH* (*MAB_3543c*) and *rshA* (*MAB_3542c*) respectively, for this investigation. We subsequently generated a *rshA* knockout strain (Δ*rhsA*) to study the role of *rshA* gene in drug resistance. The drug susceptibility difference between WT and Δ*rhsA* was not significant (data not shown). Notably, the *rshA* gene and *sigH* lie in the same cluster. Mab SigH shares 84% amino acid identity with Mtb SigH (Figure S1). The stress response factor SigH plays a crucial role in regulating responses to heat and oxidative stress and has been extensively studied in both Mtb and Msm (25). Overexpression of *sigH* in WT (OEMab*sigH*) resulted in a significant increase in resistance to TIG and FQs (Table 1).

**Table 1.**
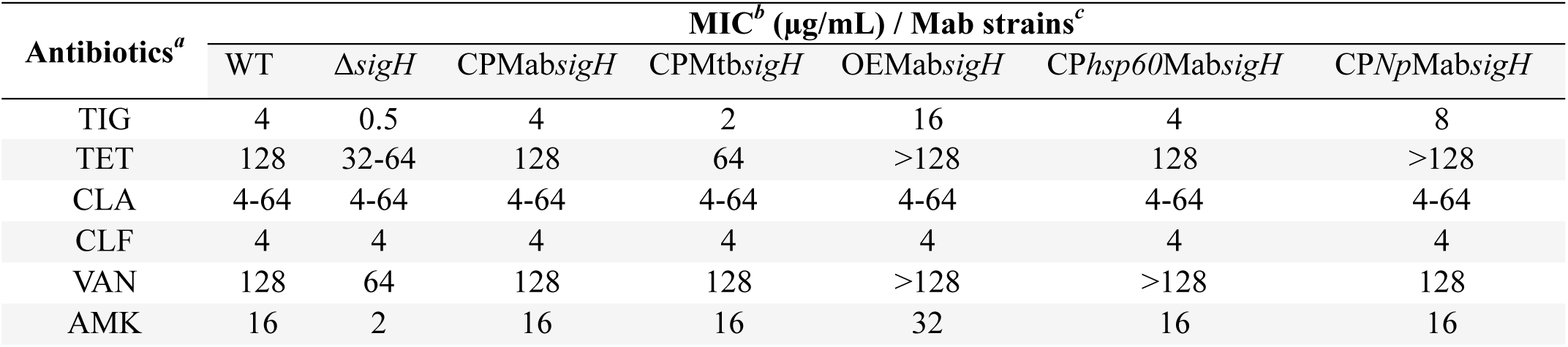
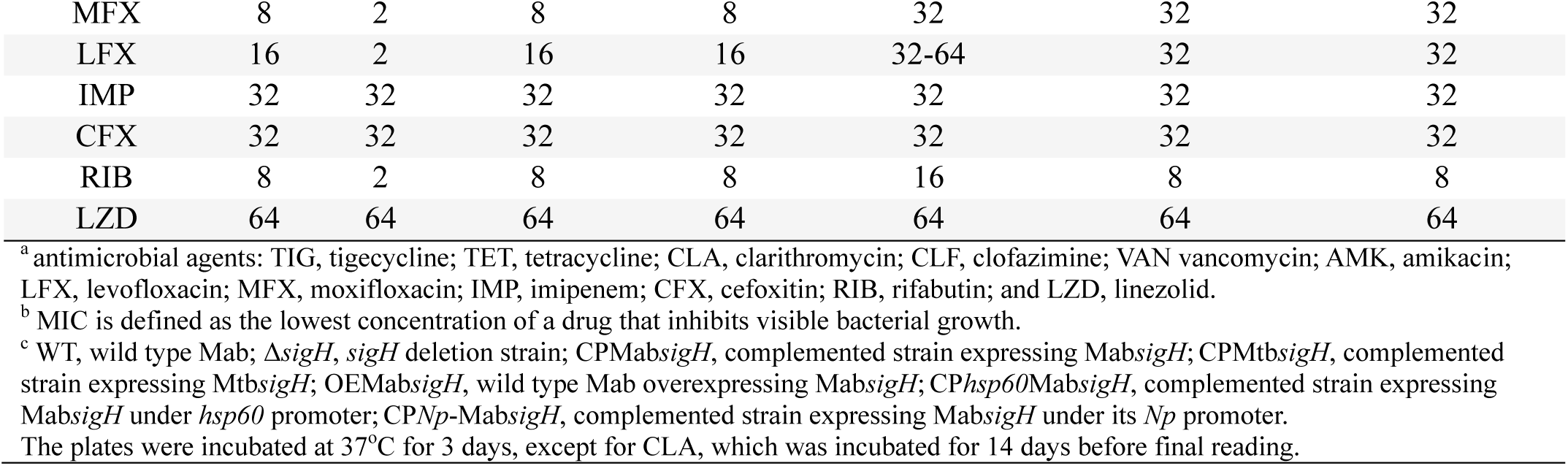
MICs of the different Mab strains.

### Deletion of *sigH* increases Mab hypersensitivity to multiple antibiotics

To investigate the role of *sigH* in multiple drug resistance, we constructed an in-frame deletion strain of Mab for *sigH* (Δ*sigH*) using CRISPR/Cpf1-assisted recombineering. The sensitivity of WT, Δ*sigH*, and its complemented strains was assessed via broth microdilution and spot growth inhibition on agar plates. Drug susceptibility testing revealed that Δ*sigH* exhibited increased sensitivity to multiple antibiotics, including ribosome-targeting agents such as TIG), TET, and AMK, as well as VAN), rifabutin RIB, and fluoroquinolones LFX and MFX. To further verify the role of *sigH* in drug resistance in Mab, complemented strains were constructed by reintroducing Mab *sigH* or its homolog from Mtb (Mtb *sigH*). The complemented strains restored resistance to the drugs (Figure 1, Table 1). Additionally, complemented strains were constructed by expressing Mab*sigH* under its native promoter (*Np*) (CP*Np*Mab*sigH*) or the strong mycobacterial *hsp60* promoter (CP*hsp60*Mab*sigH*). Complementation under either promoter led to the restoration of the drug-resistance phenotype (Table 1). These results suggest that *sigH* plays a significant role in intrinsic multiple drug resistance in Mab.

**Figure 1:**
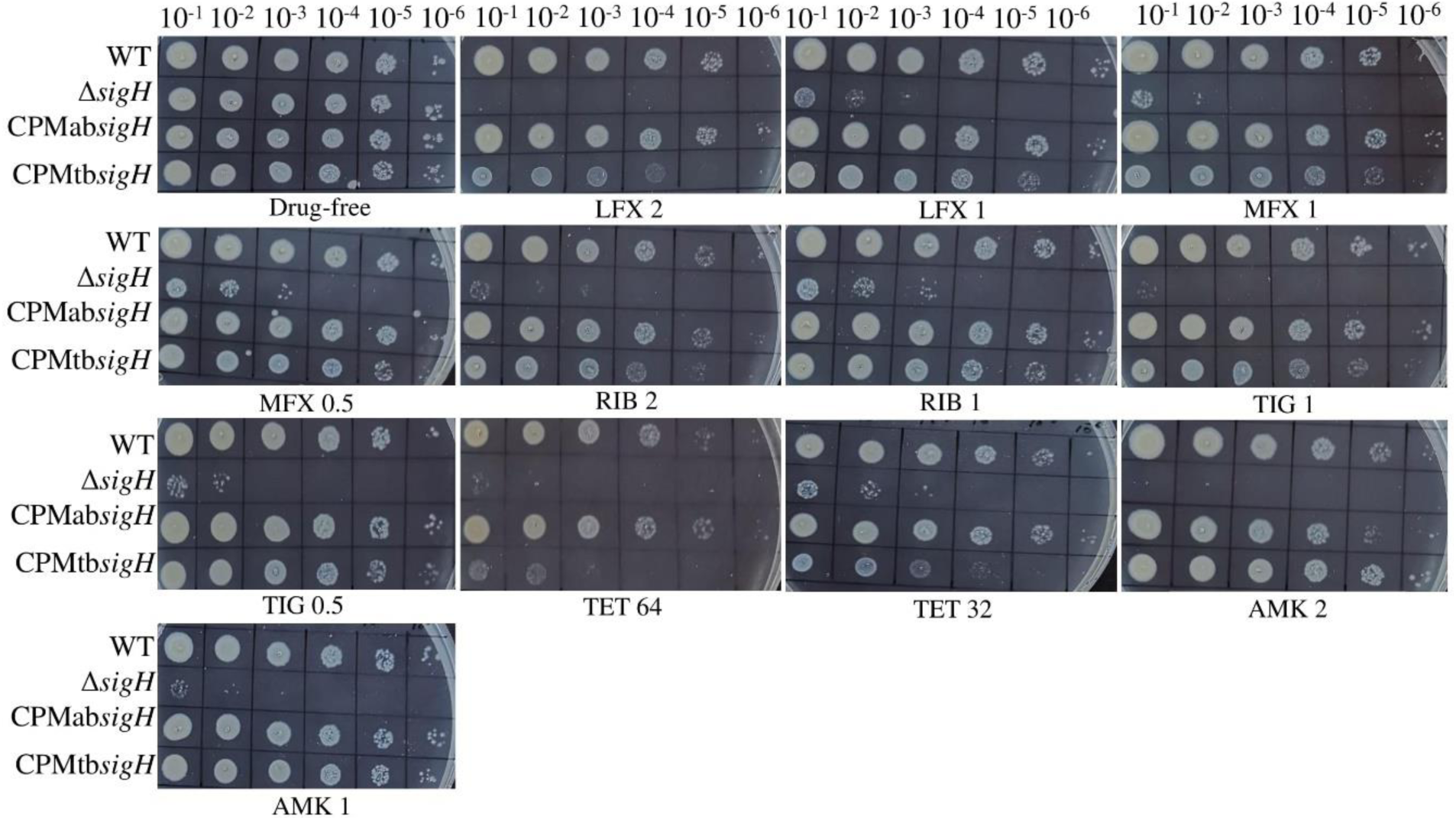
Susceptibilities of different Mab strains to different antibiotics on 7H10 agar plates. Strains were propagated in 7H9 medium at 37 °C until reaching an OD_600_ nm of 0.6. Subsequently, ten-fold serial dilutions were applied and aliquoted onto plain Middlebrook 7H10 agar (drug-free serving as a control) alongside plates supplemented with varying drug concentrations (µg/mL). Following a 3-day incubation at 37 °C, the plates were examined. TIG, tigecycline; TET, tetracycline; AMK, amikacin; LFX, levofloxacin; MFX, moxifloxacin and RIB, rifabutin.

### Deletion of *sigH* influences stress responses in Mab

SigH is a crucial regulator of a large transcriptional network that responds to heat and oxidative stress in Mtb (47). It plays an important role in virulence in animal infection models and in responding to extracellular stresses and intracellular survival (48). To investigate the role of *sigH* in Mab stress responses, we conducted heat stress assays as well as various oxidative stress assays. The diamide induction assay revealed that the Δ*sigH* strain is highly sensitive to thiol-specific oxidation by diamide. This sensitivity was evident from a larger inhibition zone in the Δ*sigH* strain (35 mm) compared to WT (20 mm) and CPMab*sigH* (26 mm) (Figure 2A; Table S1). After 8 hours, the percentages of survivors for WT, Δ*sigH*, and CPMab*sigH*, respectively, under the different conditions were as follows: Diamide (40.38%, 9.71%, and 36.25%), H₂O₂ (64.99%, 25.74%, and 60.55%), and heat (62.40%, 40.90%, and 53.26%) (Figure 2B-D).

**Figure 2:**
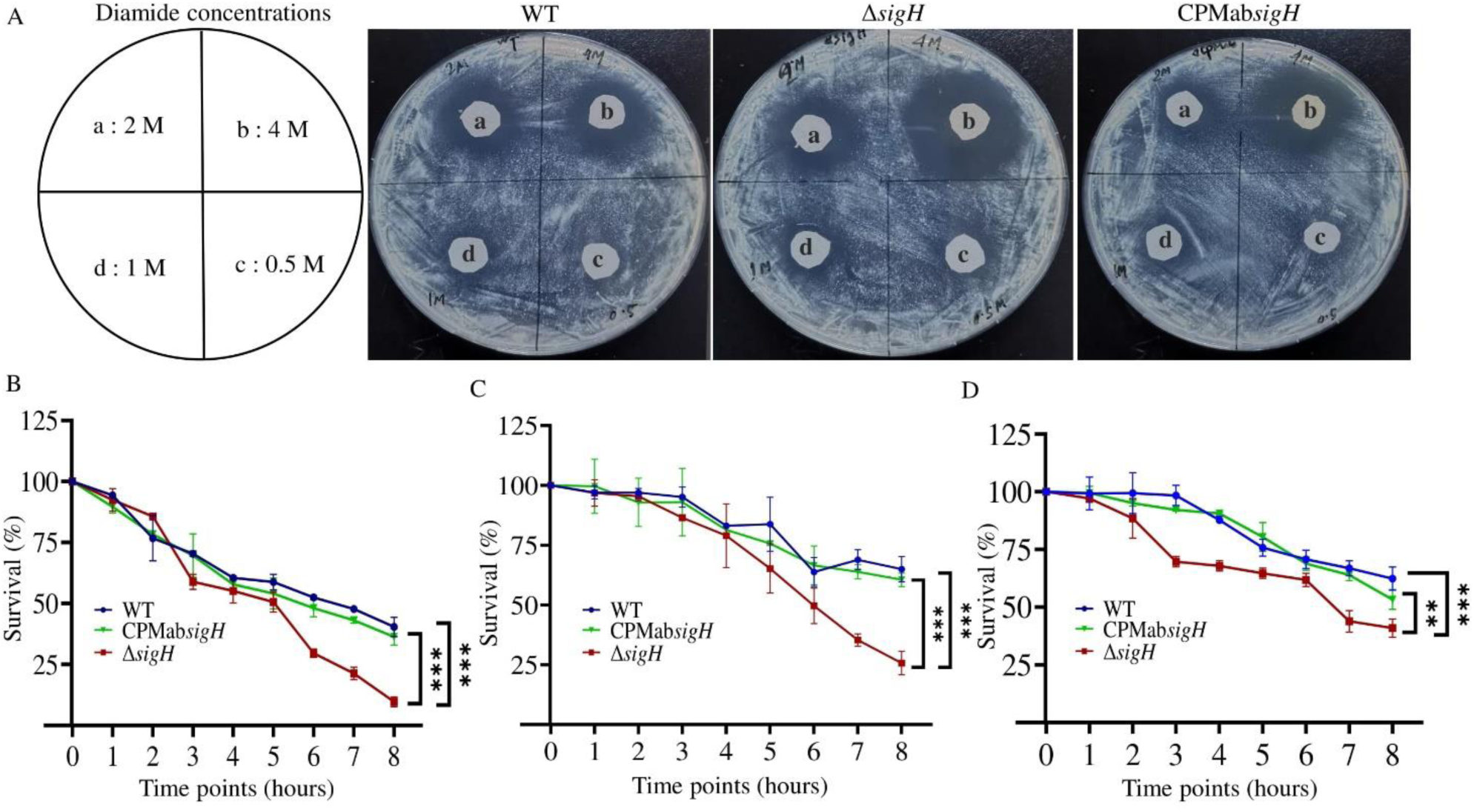
Sensitivity and survival rates of WT, Δ*sigH*, and CPMab*sigH* under stress conditions. Sensitivity of WT, Δ*sigH*, and CPMab*sigH* to different concentrations (M) of diamide (A) as well as their survival in the presence of diamide (50 mM) (B), H_2_O_2_ (50 mM) (C), and heat (45°C) (D).

### Identification of differentially expressed genes and analysis of their correlations among WT, Δ*sigH*, and CPMab*sigH*

The raw reads of transcriptome data were examined to gain insights into gene expression changes in Δ*sigH*. Several DEGs were found in both Δ*sigH* and CPMab*sigH* when compared to WT. Distance heat maps depicting the expression of all genes were utilized to hierarchically cluster the relationships among samples, thereby accurately reflecting inter-sample relationships (Figure 3A). Principal component analysis (PCA) was employed to represent the total variance and correlations among the samples (Figure 3B). This analysis revealed significant differences between Δ*sigH* and both WT and CPMab*sigH*. A correlation heat map was plotted to better understand the DEGs and relative expression patterns of shared genes among WT, CPMab*sigH*, and Δ*sigH*. The top 30 DEGs are presented in Figure 3C and supplementary files (XLS 1; XLS 2).

**Figure 3:**
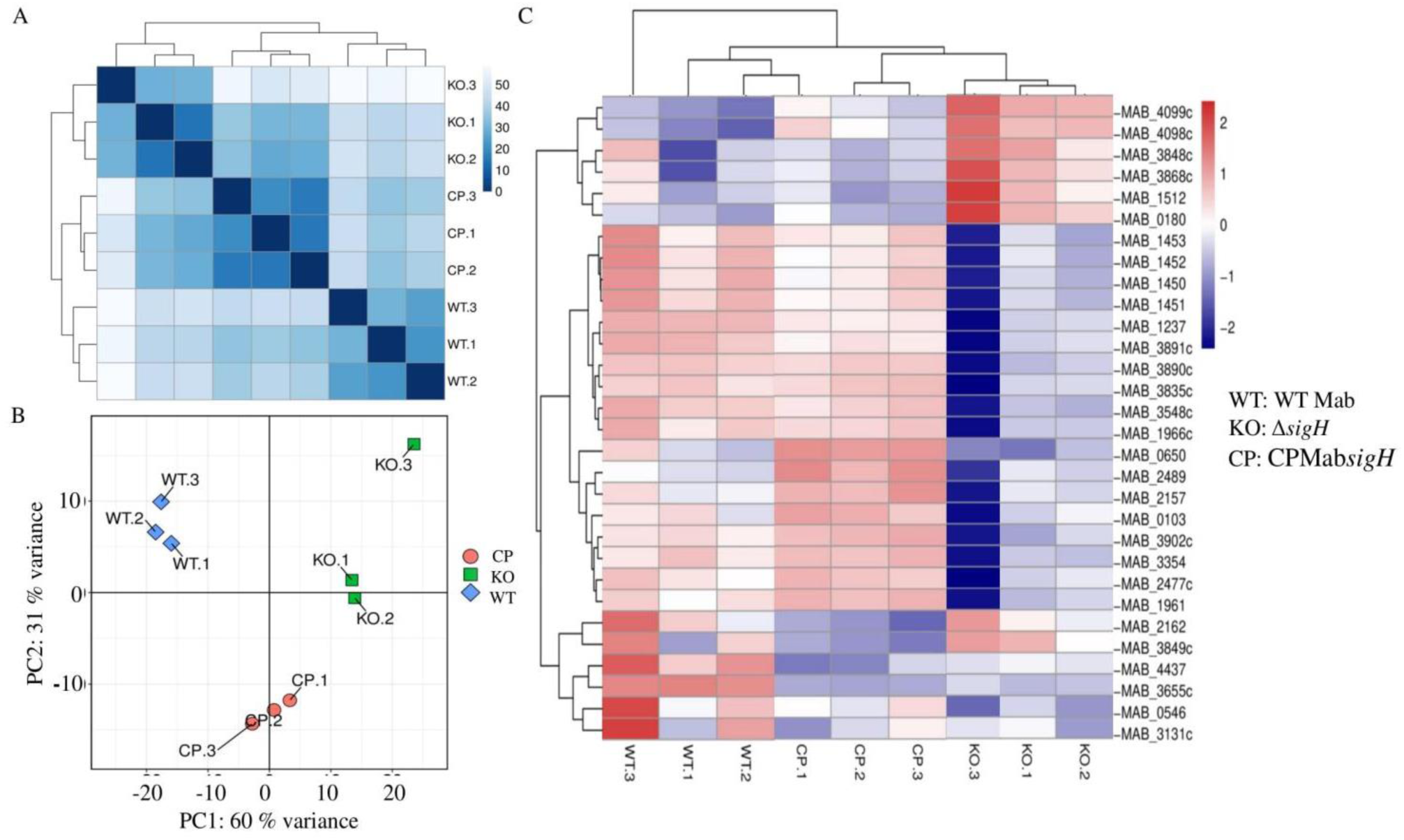
Identification of differentially expressed genes and analysis of their correlations among WT, Δ*sigH*, and CPMab*sigH*. Hierarchical clustering based on the expression of all genes can accurately reflect the relationships among samples (A). The first principal component (PC1) and the second principal component (PC2) are plotted in a two-dimensional coordinate graph, with the values in brackets on the axis labels representing the percentage of the total variance explained by each principal component (B). Based on gene expression, we performed hierarchical clustering analysis to explore the relationships between samples and genes. In the figure, each column represents a sample, and each row represents a gene. Different colors indicate the expression levels of genes across various samples. Red indicates higher expression, while blue indicates lower expression. The figure shows the top 30 gene expressions resulting from the clustering analysis (C). Note: WT: WT Mab; KO, Δ*sigH*; CP, CPMab*sigH*.

### Deletion of *sigH* affects global gene expression in Mab

To better understand the phenotype of Δ*sigH*, we identified genes affected by *sigH* deletion. Whole transcriptome analysis was used to find DEGs in Δ*sigH*. DEGs were analyzed across three comparison groups: Δ*sigH* vs. WT, WT vs. CPMab*sigH*, and Δ*sigH* vs. CPMab*sigH*. We identified 863 DEGs in the Δ*sigH* vs. WT group, with 451 upregulated and 372 downregulated. In the WT vs. CPMab*sigH* group, 464 DEGs were identified (213 upregulated, 251 downregulated). In the Δ*sigH* vs. CPMab*sigH* group, 579 DEGs were identified (353 upregulated, 226 downregulated) (Figure 4A). The highest number of DEGs was observed in the Δ*sigH* vs. WT group. Notably, twelve genes exhibited the lowest expression levels in the Δ*sigH* vs. WT group compared to the WT vs. CPMab*sigH* group (Table 2; XLS 3). These genes include *MAB_4143c*, *MAB_1362*, *MAB_4735*, *MAB_3016c*, *MAB_4694c*, *MAB_2462*, *MAB_4234c*, *MAB_4122*, *MAB_2461*, *MAB_4843*, *MAB_2459*, and *MAB_2460*, encoding putative anti-ECF sigma factor, starvation-induced DNA protecting protein/Ferritin and Dps, probable alternative RNA polymerase sigma factor, glycosyltransferase, sulfonate ABC transporter periplasmic protein, reduced flavin mononucleotide (riboflavin 5′-phosphate) (FMNH_2_) utilizing oxygenase, acyl-CoA dehydrogenase, and many conserved hypothetical proteins. A high increase in the expression of putative YrbE and MCE family proteins was observed in Δ*sigH* compared to WT (Table 2), while several putative sigma and anti-sigma factors (*MAB_3028*, *MAB_3388c*, *MAB_3016c*, *MAB_3549c*, *MAB_3548c*, *MAB_3546c*, *MAB_3542c*, *MAB_3539c*, and *MAB_3538*) were downregulated (Table S2). Other DEGs in the Δ*sigH* vs. WT group are listed in supplementary materials (XLS 3). The global differential gene expression across different comparison groups is visualized as a heatmap (Figure 4B) and a volcano plot (Figure 4C). These results suggest that SigH plays a significant role in shaping global gene expression in Mab.

**Figure 4:**
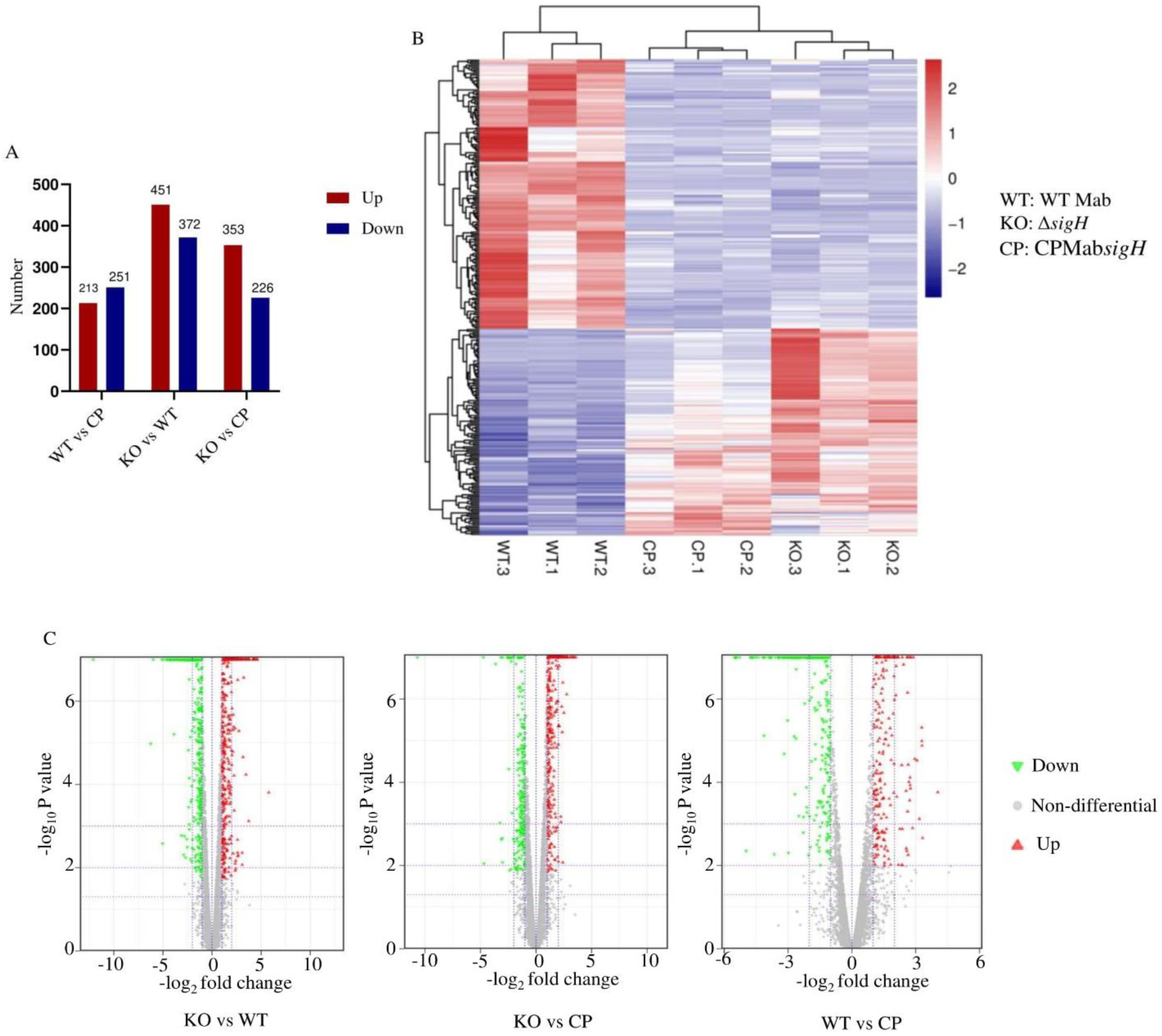
DEGs analysis of different comparison groups. The bar graph illustrates the number of upregulated and downregulated DEGs (A). The heat map displays DEGs across different comparison groups; the color intensity indicates the relative expression level of the genes (B). The volcano plot presents differentially expressed genes in the comparison groups, with each point representing a gene. The horizontal axis denotes the log_2_ fold change, while the vertical axis represents the negative log_10_ of the p-value. Red dots indicate upregulated DEGs, green dots signify downregulated DEGs (down-regulated on the left and up-regulated on the right), and gray dots represent non-differentially expressed genes (C). Only genes with |log_2_FoldChange| > 0 and *p <* 0.05 are included in the volcano plot. Note: WT: WT Mab; KO, Δ*sigH*; CP, CPMab*sigH*.

**Table 2:**
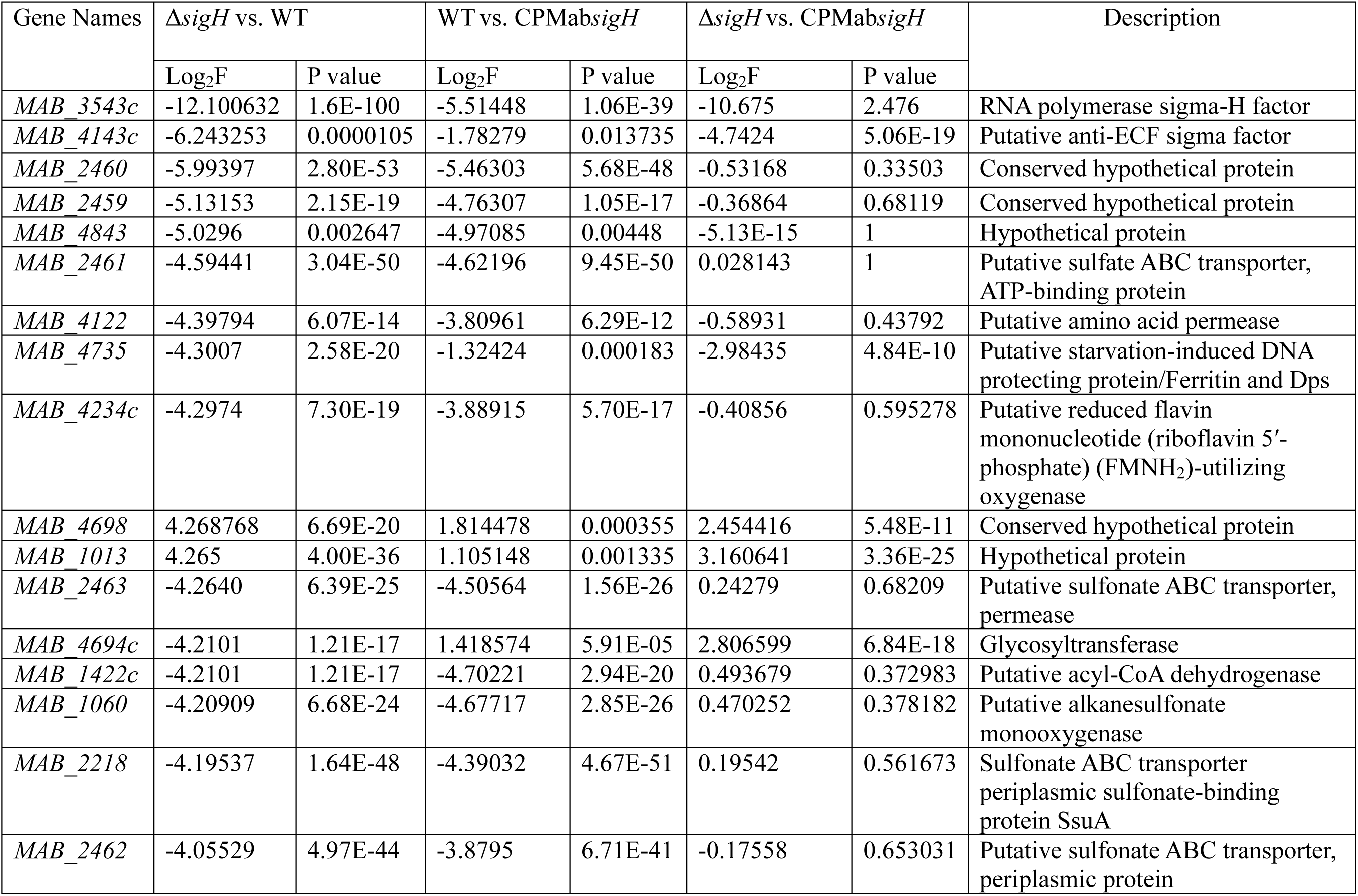
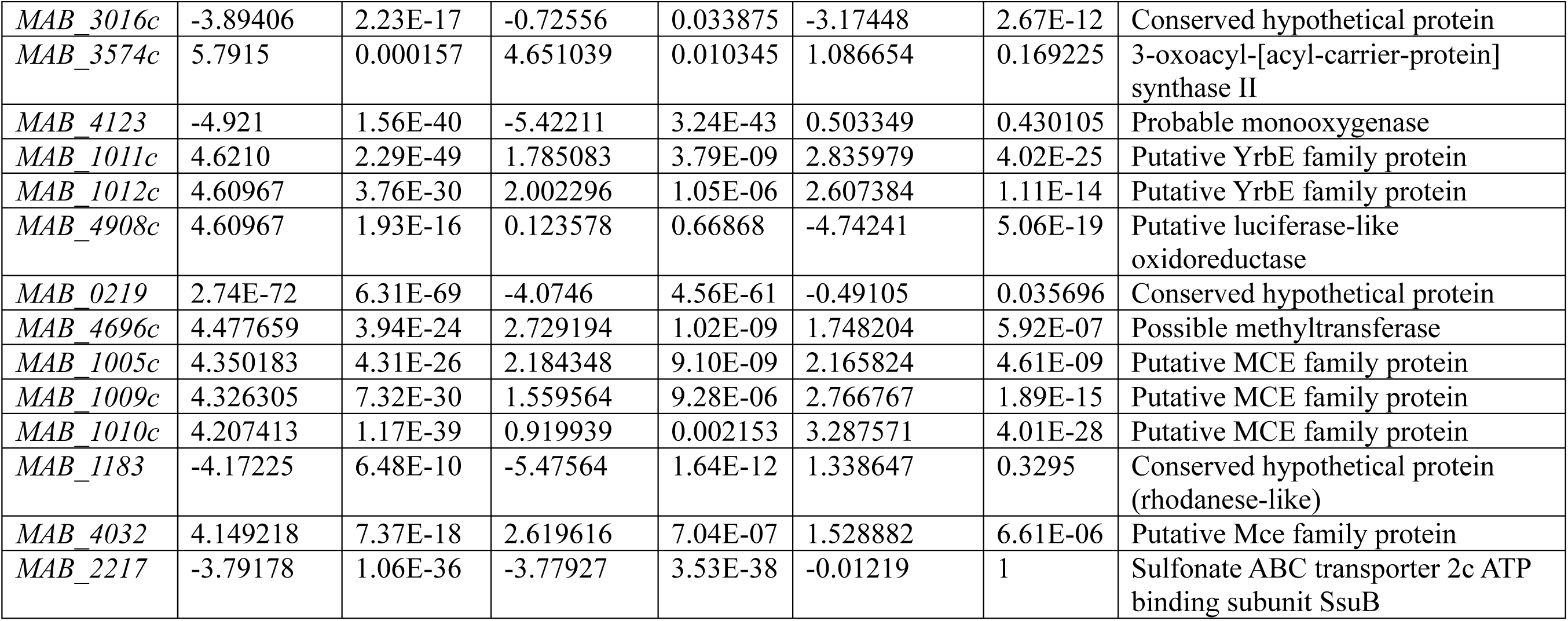
The most significant alterations in the transcriptome of DEGs across various Mab groups identified using log_2_ fold change values.

Following the change in global gene expression after *sigH* deletion, we performed KEGG and GO gene enrichment analyses to determine enriched pathways in Δ*sigH*. The enriched pathways included those involved in metabolism, cellular processes, human diseases, and the processing of genetic and environmental information (XLS 4; XLS 5). The distribution of KEGG DEGs across the different comparison groups is as follows: Δ*sigH* vs. WT (45 upregulated, 76 downregulated); Δ*sigH* vs. CPMab*sigH* (57 upregulated, 10 downregulated); and WT vs. CPMab*sigH* (45 upregulated, 76 downregulated). The enriched pathways are detected in Δ*sigH* vs. WT (85 enriched pathways), WT vs. CPMab*sigH* (64 enriched pathways), and Δ*sigH* vs. CPMab*sigH* (72 enriched pathways) groups (Figure S6; XLS 6). For the KEGG enrichment analysis of DEGs, the DEGs in the Δ*sigH* vs. WT group were mainly enriched in ABC-type transporters. Further, GO enrichment analysis identified changes in DEGs associated with sigma factor, transporter, cofactor, and transmembrane transporter activities (Figure S6; XLS 5). These results emphasize that *sigH* inactivation has a profound impact on transcriptional, sigma, anti-sigma, oxidative, and ABC transporter-related gene functions.

### SigH is involved in drug resistance perhaps by influencing other drug resistance determinants in Mab

SigH is implicated in drug resistance by influencing other drug resistance determinants in Mab. The multi-drug resistance WhiB7 master regulator induces the transcription of *sigH*. SigH is an alternative sigma factor associated with the transcriptional regulation of genes responsible for mycobacterial stress responses, including oxidative, heat, and nitrosative stress. It is negatively regulated by *rshA* (15). This sigma factor controls visible physiological alterations and modifies gene expression patterns during antibiotic treatments and diverse environmental stresses, playing a significant role in pathogen drug resistance (49, 50, 51). In this study, several genes were found to be down-regulated in Δ*sigH* compared to WT and CPMab*sigH* strains. Examples include *MAB_1362*, *MAB_4143c*, *MAB_3028*, *MAB_3388c*, *MAB_3542c*, and *MAB_3016c*, all previously associated with drug resistance in Mab. For instance, under *sigH* regulation, *MAB_1362* influences intrinsic resistance to AMK, streptomycin (STR), and apramycin (APR) (52). Additionally, the deletion of *MAB_3542c* increases Mab’s sensitivity to TIG (53), while a mutation in *MAB_3388c* (*serB2*) has been linked to cross-tolerance to CFX and MFX (54). Thus, our results suggest that SigH influences drug resistance in Mab by modulating the expression of other genes directly involved in drug resistance.

### Deletion of *sigH* enhanced Mab cell wall permeability

The deletion of *sigH* affects the cell wall permeability of Mab. The hydrophilic fluorescent dye ethidium bromide (EtBr) can intercalate into DNA and RNA and penetrate the cell walls of mycobacteria. Mycobacterial intrinsic resistance to several antimicrobials is thought to be mainly due to decreased cell wall permeability and active efflux mechanisms (55). To investigate whether the *sigH* deletion affects the cell wall permeability of Mab, we performed an EtBr accumulation assay. We observed that Δ*sigH* accumulated more EtBr relative to WT, and complementation with CPMab*sigH* restored the phenotype partially (Figure 5). These findings suggest a possible increase in cell envelope permeability following *sigH* deletion, highlighting the significance of *sigH* in maintaining Mab cell wall integrity.

**Figure 5:**
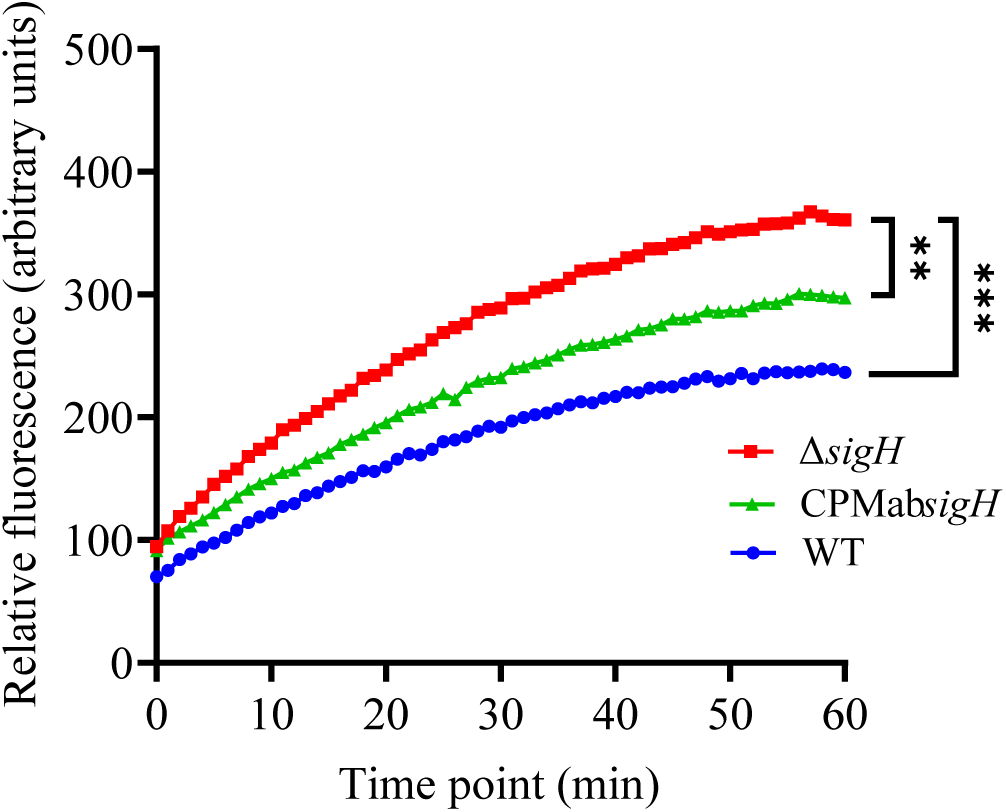
Estimation of cell envelope permeability. Accumulation of EtBr in the cells of WT, Δ*sigH,* and CPMab*sigH* strains.

## Discussion

The rapidly growing NTM species Mab is an emerging healthcare-associated opportunistic pathogen characterized by its drug-resistant phenotype and high morbidity and mortality rates (56, 57). Mab causes chronic pulmonary diseases that are difficult to manage due to inherent drug resistance, posing a significant public health threat. This underscores the need for investigations to identify novel therapeutic targets and effective treatment options.

Bacterial pathogens regulate their gene expression in response to environmental cues during infection, which is crucial for virulence (58). SigH, a sigma factor, plays a critical role in regulating transcription and stress responses in Mtb, Msm, and *Mycobacterium avium ssp. paratuberculosis* (Mav). SigH controls various genes, including other extracytoplasmic function sigma factors and redox systems (23, 59). It is particularly important for the pathogen’s response to heat and oxidative stress, roles extensively studied in Mtb and Msm. Although less well-characterized in Mab, SigH has been linked to resistance against TIG and AMK (52, 60). Notably, the peptide sequence of Mab SigH shares 84% similarity with that of Mtb SigH. In this study, we demonstrated that SigH confers resistance not only to TIG and AMK but also to multiple drugs in Mab (Table 1, Figure 1). The increased sensitivity of the Δ*sigH* strain to antibiotics such as LFX, MFX, TIG, TET, AMK, VAN, and RIB highlights this role. Complementation of *sigH* in Δ*sigH* restored drug resistance, suggesting that SigH’s role in drug resistance is conserved between Mab and Mtb (16, 17, 61). Consistently, Δ*sigH* showed heightened sensitivity to stressors like diamide, H₂O₂, and heat, significantly affecting its survival (Figure 2). These findings indicate that SigH’s role in stress response is conserved across these mycobacterial species.

Sigma factors are global gene regulators and key transcription activators in mycobacterial pathogenesis. They bind to RNA polymerase, enhancing affinity for specific promoters (62). To investigate *sigH’s* influence on global gene expression in Mab, we performed transcriptomic profiling of WT, Δ*sigH*, and complemented CPMabΔ*sigH*. Significant changes were observed in the gene expression profile of Δ*sigH* compared to WT, while CPMab*sigH* closely resembled WT, indicating partial restoration of SigH-associated regulation (Figure 3). A large set of genes was differentially expressed: 863 DEGs (451 upregulated, 372 downregulated) in Δ*sigH* vs. WT; 464 (213 upregulated, 251 downregulated) in WT vs. CPMab*sigH*; and 579 (353 upregulated, 226 downregulated) in Δ*sigH* vs. CPMab*sigH* (Figure 4, Table 2). KEGG enrichment analysis revealed that the most downregulated (log_2_FC ≤ −3) genes in Δ*sigH* were ABC-type transporters (22%), potentially explaining the increased antibiotic sensitivity. ABC-type transporters facilitate import/export and drug efflux in mycobacteria (63, 64, 65), so disruption of *sigH* likely impairs these functions, reducing drug tolerance (Table 1, Figure 1). This is supported by EtBr accumulation in Δ*sigH* (Figure 5). Additionally, the downregulation of genes associated with antibiotic resistance was observed, including *MAB_1362* (alternative RNA polymerase sigma factor), *MAB_3542c* (anti-sigma factor), and *MAB_3388c* (phosphoserine phosphatase *serB2*) (52, 53, 54). Although we did not test sensitivity to STR and APR, these results align with the observed sensitivity profile. Interestingly, 12 genes were highly upregulated (log_2_FC ≥ 2), with five genes (two *yrbE* and three *mce* genes) within an *mce* operon. MCE proteins are associated with virulence and stress responses in mycobacteria (66). Previous studies suggest that oxidative stress, hypoxia, and nutrient deprivation can modify MCE protein expression, implying a role in stress responses (67, 68, 69). Another *mce* gene (*MAB_4032*) was also highly upregulated. We speculate that following *sigH* disruption and subsequent downregulation of antibiotic resistance genes, Mab attempted to reinforce its cell wall and counter external stressors by upregulating *mce* genes. However, further experimentation is needed to validate this hypothesis.

A synthetic drug-like molecule, SMARt-420 (Small molecule aborting resistance), has been shown to inhibit the DNA binding effect of EthR2 (Rv0078), which is a transcriptional repressor of EthA2 in Mtb. By doing so, SMARt-420 profoundly facilitates the bioactivation and antibacterial effect of ethionamide against Mtb (70). If inhibition of the transcriptional regulator for a single gene would have this profound impact, it would be reasonable to suspect that a protein with genome-wide transcriptional regulatory effect could be an ideal target for drug development. In fact, SigH could be an ideal target for drug development against Mab due to its obvious impact on overall gene expression in this pathogen, as this would have great impact on several pathways that could be involved in drug resistance and stress response.

## Summary

We have identified the role of the mycobacterial sigma factor SigH in stress responses and multi-drug resistance in Mab. We hypothesize that disrupting *sigH* in Mab leads to the downregulation of multiple drug resistance determinants, thereby increasing bacterial sensitivity to drugs. Additionally, we suggest that as a compensatory mechanism for this downregulation, the bacteria upregulate the expression of certain proteins, such as YrbE and MCE family proteins. Although these findings require further experimental validation, this study lays the foundation for future mechanistic studies to establish the potential of SigH as a drug target in Mab.

## Author contributions

**Md Shah Alam**: Conceptualization (equal); investigation (lead); methodology (lead); validation (equal); visualization (lead); formal analysis (lead); writing— original draft (lead); writing— review and editing (equal); data curation (lead). **Mst Sumaia Khatun:** Methodology (equal); investigation (equal); formal analysis (equal) and writing— original draft (supporting). **Buhari Yusuf**: Original draft (supporting) and writing— review (lead). **Lijie Li**: Data curation (equal) and formal analysis (supporting). **Aweke Mulu Belachew**: Formal analysis (supporting); software (supporting). **Haftay Abraha Tadesse**: Formal analysis (supporting); software (supporting). **Jingran Zhang**: Original draft (supporting) and formal analysis (supporting). **Xirong Tian**: Formal analysis (supporting); validation (supporting). **Cuiting Fang**: Formal analysis (supporting); validation (supporting). **Yamin Gao**: Formal analysis (supporting); validation (supporting). **Zhiyong Liu**: Formal analysis (supporting); validation (supporting). **H.M. Adnan Hameed**: writing—original draft (supporting) and software (supporting). **Jinxing Hu**: Resources (supporting), writing—review, and editing (supporting). Xinwen Chen: Resources (supporting), writing—review, and editing (supporting). **Nanshan Zhong**: Validation (supporting), and writing—review. **Shuai Wang**: Conceptualization (equal); project administration (lead); funding acquisition (lead); resources (supporting); validation (equal) and writing—review and editing (equal); **Tianyu Zhang**: Conceptualization (equal); project administration (lead); funding acquisition (lead); resources (supporting), validation (equal) and writing—review and editing (equal).

## Acknowledgments

This work was supported by the National Key R&D Program of China (2021YFA1300904, 2023YFF0713605), the National Natural Science Foundation of China (32300152), partially by Guangdong Provincial Basic and Applied Basic Research Fund (2024A1515012412, 2022A1515110505), the State Key Laboratory of Respiratory Disease, Guangzhou Institute of Respiratory Diseases, First Affiliated Hospital of Guangzhou Medical University (SKLRD-Z- 202412, SKLRD-Z-202414, SKLRD-Z-202301). The founders had no role in study design, data collection, and analysis, decision to publish, or preparation of the manuscript. We also acknowledge the group of Yicheng Sun from the Institute of Pathogenic Biology Chinese Academy of Medical Sciences, for kindly sending the pJV53-Cpfl and pCR-ZEO plasmids as tools for gene deletion.

